# The development of spatiotemporal organization of episodic memory in children and its disruption in Williams Syndrome

**DOI:** 10.1101/534743

**Authors:** Marilina Mastrogiuseppe, Natasha Bertelsen, Maria Francesca Bedeschi, Sang Ah Lee

**Author notes:** **Corresponding author:** Sang Ah Lee, Department of Bio and Brain Engineering, Korea Advanced Institute of Science and Technology, Daehak-ro 291, Daejeon, 34141, Republic of Korea, Email address.

## Abstract

Recent theories of episodic memory propose that the hippocampus provides the spatiotemporal framework for episodic memories. If this is true, does the development of episodic memory depend on the binding of space and time? And does this rely, at least partly, on normal hippocampal function? We investigated the development of episodic memory in children 2–8 years of age (Study 1) and its impairment in Williams Syndrome (Study 2) by implementing a nonverbal object-placement task that dissociates the *what*, *where*, and *when* components of episodic memory. Our results indicate that the binding of space and time in memory emerges first in development around the age of 3 and is impaired in Williams Syndrome. Space-time binding both preceded and predicted success in full episodic memory (*what+where+when*), and associating objects to spatial location seemed to mediate this developmental process. Importantly, these effects were not explained by improvements in object or location memory.

## Introduction

Episodic memory (EM), the memory for personally experienced events, is set in a particular context and contains information about the time and place in which the events took place. Each experience that we live is made out of different elements, such as the content of the memory (“what”), its spatial location (“where”) and its temporal information (“when”). To create an EM representation, it is not enough to remember the single elements of an event; instead, they have to be remembered as a binded whole.

It is well known that the hippocampal formation (HF) is crucial for memory binding processes, as confirmed by several studies of patients with hippocampal damage (Elward et al., 2018; Gold et al., 2006; Baddeley, Vargha-Khadem, & Mishkin, 2001; Brizzolara et al., 2003; Hurley, Maguire, & Vargha-Khadem, 2011; Rosenbaum et al., 2011). Furthermore, although the HF has long been regarded as critical for spatial representations (O’Keefe & Nadel, 1978; Miller et al., 2013; Doeller, Barry & Burgess, 2010; O’Keefe & Dostrovsky, 1971; Tolman, 1948; Hartley et al., 2014; Killian, Jutras & Buffalo, 2012), there is now a large consensus that is also involved in the temporal organization of memories (Thavabalasingama, O’Neil & Lee, 2018; Howard & Eichenbaum, 2015; Komorowski, Manns, & Eichenbaum, 2009; McKenzie et al., 2014; Hsieh et al., 2014; Kumaran and Maguire, 2006; Lehn et al., 2009; Tubridy and Davachi, 2011).

As humans and animals move through their surroundings, memory for subsequently visited locations contributes to create a representation of space that can be used to navigate. In the same way, episodic memories are created while individuals move in their environments and interact with different objects and people in specific spatiotemporal contexts. Recent findings illustrating the properties of hippocampal and entorhinal cells involved in the representation of episodic time (Rubin et al., 2015; Cai, D. J. et al., 2016; Mau et al., 2018; Eichenbaum, 2014; MacDonald et al., 2011; Tsao et al., 2018), as well as functional imaging studies on patients with hippocampal damages showing specific impairment in the temporal organization of memories (e.g. Konkel, Warren, Duff, Tranel & Cohen, 2008; Spiers, Burgess, Hartley, Vargha-Khadem, O’Keefe J., 2001), provide considerable support for the HF as representing not only spatial but also temporal information.

Given this perspective on hippocampal function, it has been suggested that HF uses space and time as a primary scaffold for encoding and retrieving episodic memories and that other dimensions can be incorporated into this initial spatiotemporal framework if they are relevant to defining the event (Ekstrom & Ranganath, 2017). Perhaps, space and time are used by the HF to break up experiences into specific contexts, and to organize multimodal input associated with each context. In this paper we explored whether there is an ontogenetic origin of this space-time scaffold upon which EM may be constructed. For this purpose, we investigated the emergence of individual binding processes across development (Study 1) using a nonverbal object-hiding task which allowed us to dissociate the conceptual content of memories (e.g., what) from their spatial (where) and temporal (when) organization (Figure 1). If there is an ontogenetic basis to the spatiotemporal scaffold, it follows that the ability to bind space and time (where+when) should develop before, and predic the emergence of, the ability to represent a full episode (what+where+when). To further assess the role of a spacetime scaffold in EM, we also investigated how hippocampal abnormalities affect memory (Study 2) by conducting a similar test in subjects with Williams Syndrome (WS), a genetic neurodevelopmental disorder characterized by an abnormal maturation of the HF and a severe deficit in spatial cognition. If the HF is critical to the spatiotemporal binding of EM, it follows that in WS we should see a particular disruption of combining “where+when” that impairs their EM representation, that constrast with a relatively spared object-related (“what”) memory.

**Figure 1.**
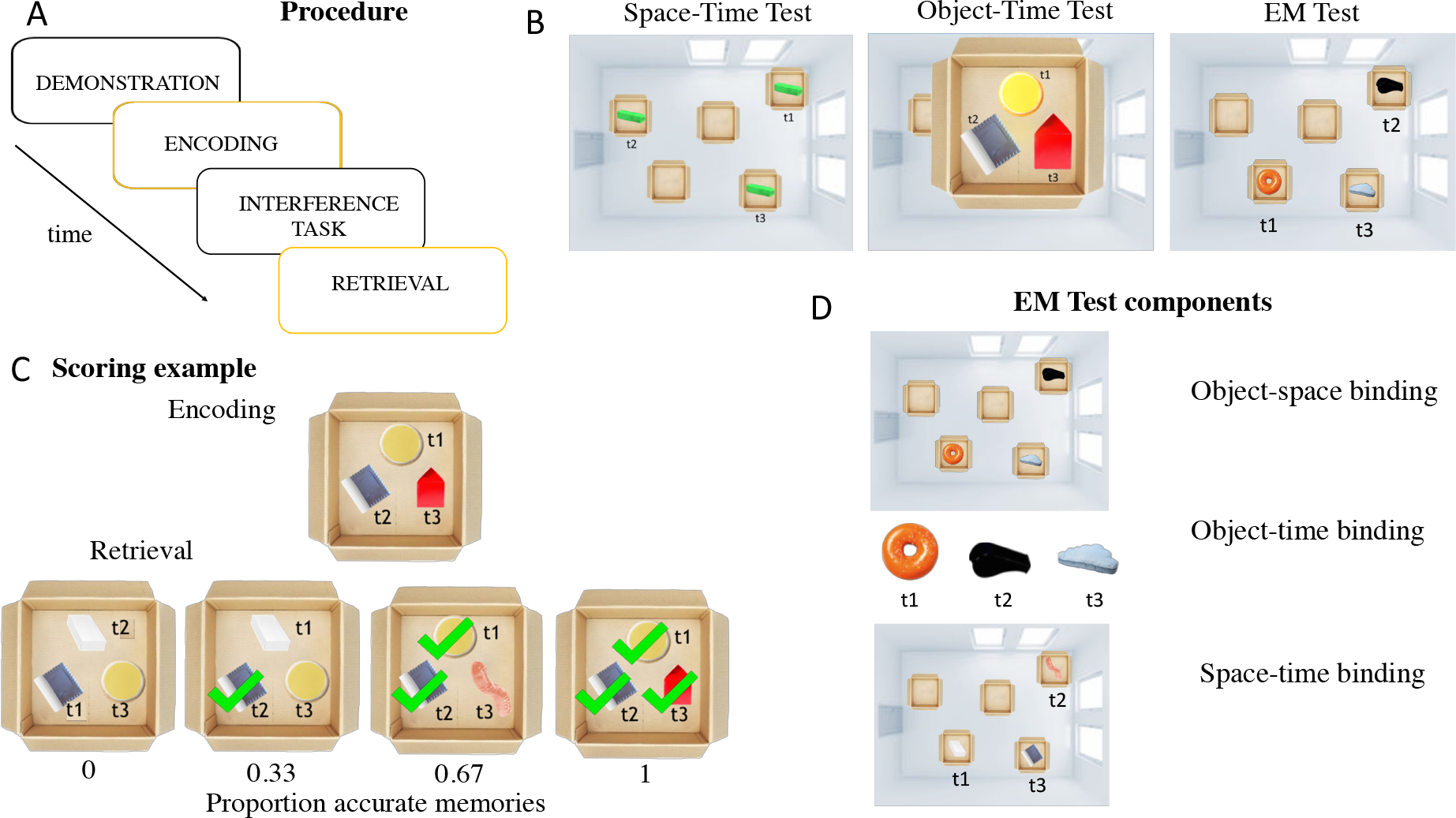
Episodic Memory Task. **A.**The experimenter sequentially placed three objects in boxes laid out in the experimental room while the participant was watching (demonstration phase), and then the participant was asked to copy the experimenter by placing the objects themselves (encoding phase). Finally, after a delay (verbal interference task), the participant was asked to re-enact his own previous actions (retrieval phase). **B.** Each experimental session was composed of three tests: (i) The Space-Time Test required the combination of location and temporal sequence information but kept the objects constant; (ii) The Object-Time Test involved different objects but held the spatial location constant; (iii) The EM Test required the binding of information concerning an object’s identity, a spatial location and temporal sequence into one representation. **C.** By comparing the subject’s behavior between the encoding and retrieval phases, we calculated the extent to which participants remember the single elements (object identity or spatial location) and their ability to bind them in a temporal sequence (object-time; space-time; EM). **D**. By separately analzing the binding components in the EM Test it was possible to further investigate the effect of object-space, object-time and space-time binding on EM representation.

### Study 1: Typical Development of EM binding components

Previous studies have reported that episodic memory in children develops around 6 years of age, when a mature hippocampus begins integrating the spatial and temporal context of an experience into the representation of a single event (Bruce, Dolan, & Phillips-Grant, 2000; Gogtay et al., 2006; DeMaster et al., 2013). The process of memory binding may occur even before the age of six, but in a considerably less reliable fashion (Olson & Newcombe, 2014). Studies have shown that 2-to-6 years old children have difficulty in remembering the context in which an item is learned (source memory) and are susceptible to false alarms when they see familiar items and locations rearranged and combined in unfamiliar ways (Sluzenski, Newcombe, & Ottinger, 2004; Sluzenski, Newcombe, & Kovacs, 2006; Lloyd, Doydum, & Newcombe, 2009). Most studies on EM binding processes have focused on their maturation in older children in comparison to memory abilities in adults but not on their initial emergence in early childhood, making it unclear what is the exact developmental trajectory of the episodic memory-binding process. This may be partly due to the limitations of the commonly used EM paradigms which often rely on designs that preclude their use with younger or delayed children, such as complex tasks with elaborate instructions, computer-based tasks, or verbal tasks that confound verbal skills with memory abilities (Hayne & Imuta, 2011; Simmock & Hayne, 2003). Moreover, most studies have utilized designs that exclude the component of space intended as a physical location of a moving observer, therefore limiting the elements of navigation and any potential assessment of the exact contribution of spatial information in episodic recall. However, in real life episodic memories are created in action, as we navigate our environment and interact with people and objects across space and time.

To overcome the limitations of past studies, we created a simple nonverbal task, involving active movement of the participant in 3D space (rather than just observation) to induce true real-life episodic memories. In our task, children observed an experimenter hiding various objects around the room (demonstration phase) and are invited to imitate the actions themselves (encoding phase). Then, after a verbal-naming interference task, children were asked to re-enact their own actions (retrieval phase). In order to investigate which types of memory binding are fundamental to episodic memories, the session consisted of three separate tests that allowed us to assess not only the EM representation (what+where+when) but also to isolate the specific binding of memory components with time: space-time binding (where+when), and object-time binding (what+when). We also separately analyzed the following memory binding in the EM Test: space-time, object-time, and object-space (what+where) binding (see Figure 1). We analyzed the proportions of accurately bounded object identities and/or locations across time by focusing on potential changes between the encoding and retrieval phases to observe how much of the original event has been committed to memory.

## Results

### Age-related differences in binding components across TD

Figure 2 shows performance in the three tasks across development. To assess developmental change, we split the subjects into three age groups: 2–4, 4–6, 6–8 years old.

**Figure 2.**
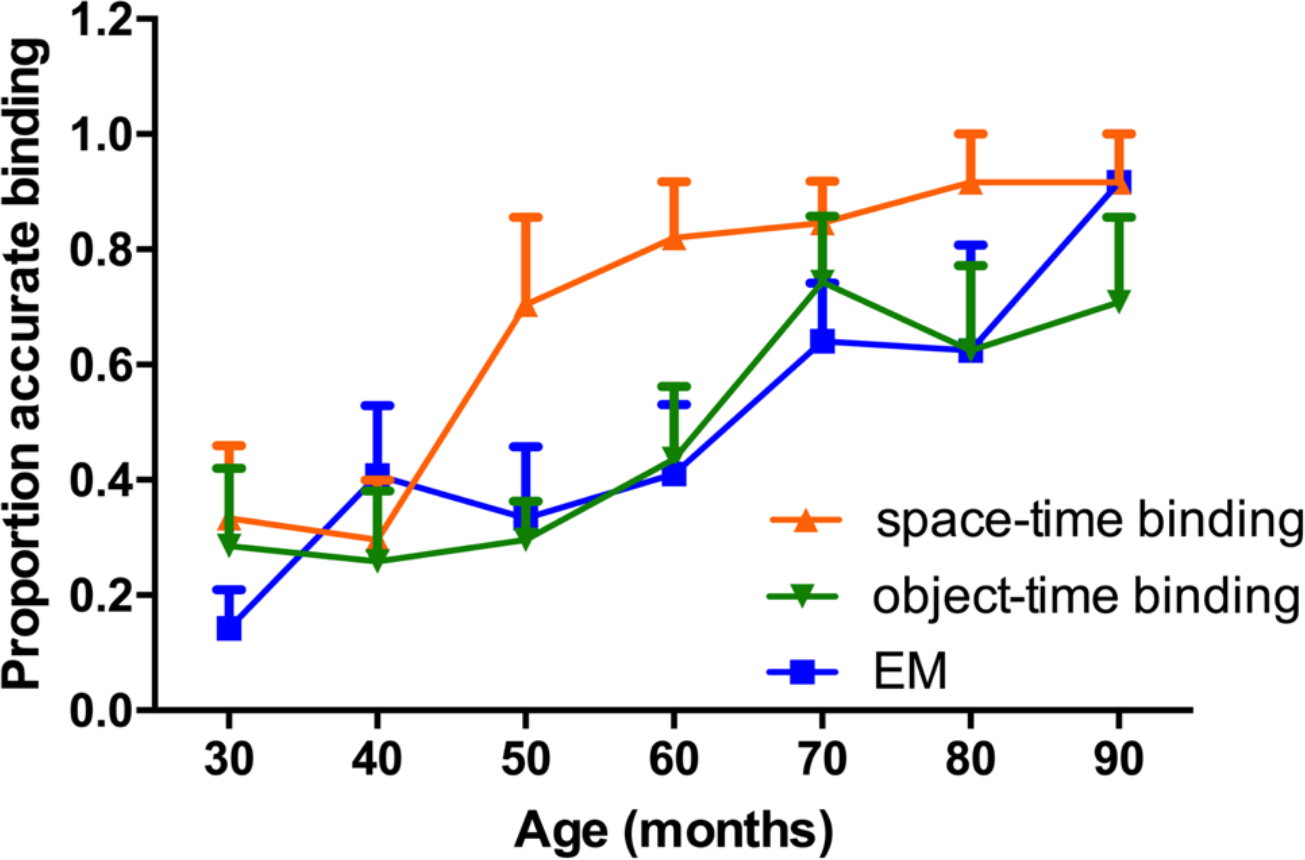
The graph presents the proportion of accurate memory binding across 2 to 8 years of age for *space-time* (where and when), *object-time* (what and when), *EM* (what, where and when). Space-time binding emerges first, followed by the other two conditions.

Performance on all three tasks varied significantly by age-group (Kruskal Wallis one-way ANOVA (Figure 3 A) H _(2)_ =12.278, p=.002; H _(2)_ =23.575, p=.000008; H _(2)_ =12.178, p=.002, respectively). 2–4-year-olds and 4–6-year-olds differed significantly only in Space-Time Test (H _(2)_ =-20.042, p=.001, Bonferroni-corrected). In the other two tests, 2–4- and 4–6-year olds performed equally well and were both significantly worse compared to the oldest group of children aged 6–8 (Object-Time Test: 2–4 years: H _(2)_ = −18.934, p=.004; 4–6 years: H _(2)_ = −13.278, p=.031; EM Test: 2–4 years: H _(2)_ = −18.282, p=.005, 4–6 years: H _(2)_ = −14.074, p=.020). These results suggest an earlier maturation of space-time binding between 2–4 years of age, followed by a protracted development of object-time binding, and EM. These patterns of results cannot be attributed simply to a memory of motor movements. In fact, a subset of subjects were presented with an obstacle at the retrieval phase that forced them to take a roundabout path that was different from the encoding phase (gated condition), and no differences in performance were founded between gated vs normal condition (F_(1,8)_ = 4.143, p >.05).

**Figure 3.**
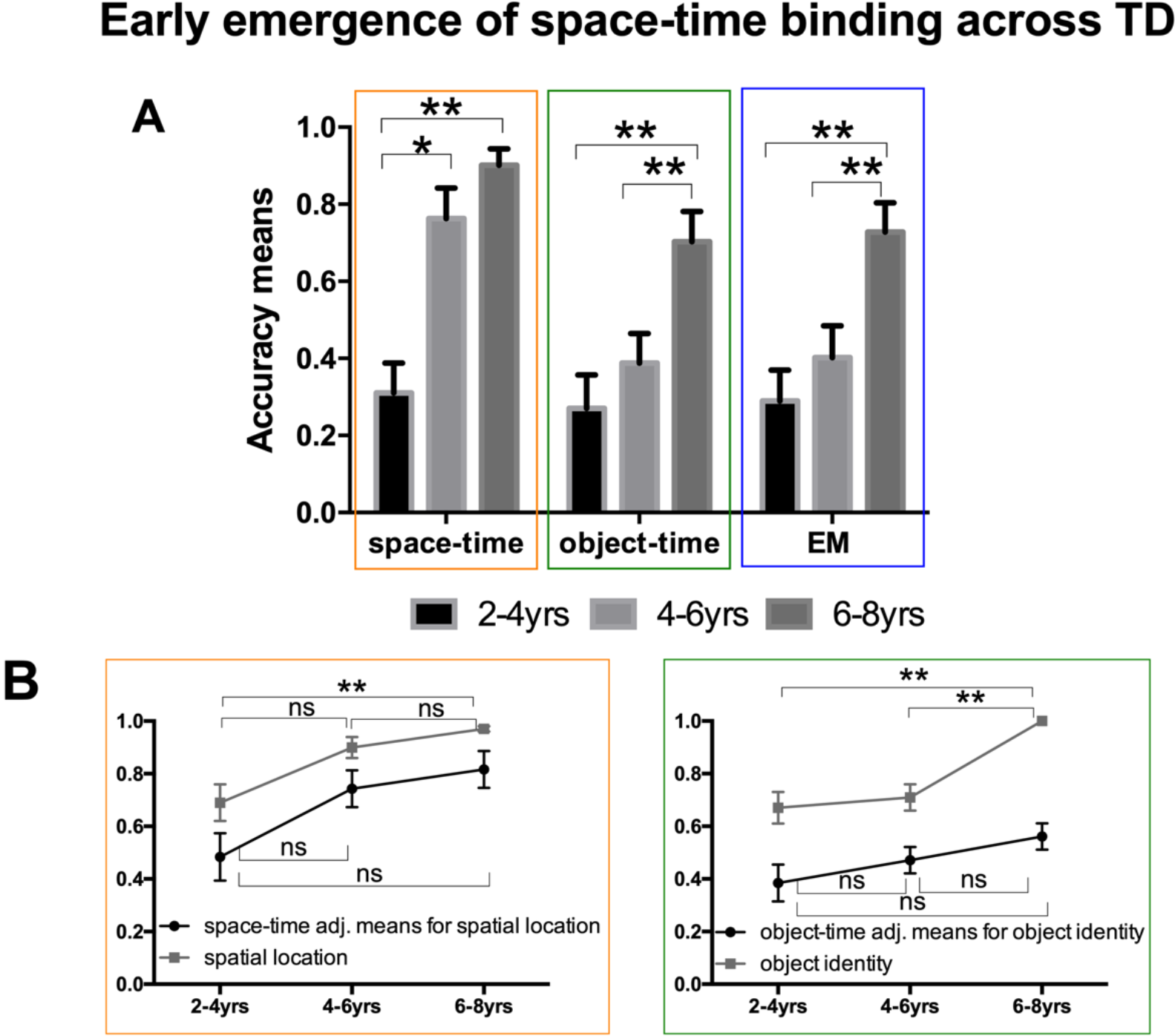
Typical development of binding components of EM *A.* The graphs present the accuracy means for Space-Time, Object-Time and EM Tests in the TD sample divided by age-groups (2–4; 4–6; 6–8 yrs). *B.* The graphs present the means for spatial location accuracy, and space-time binding adjusted for spatial location accuracy (left), and for object identity accuracy, and object-time binding adjusted for object identity accuracy (right). Adjusted means were calculated using GZLM. ns; p > 0.05 *; p ≤ 0.05 **

Interstingly, in normal condition spatial location accuracy showed a significant difference across age groups (H _(2)_ = 16.835, p=.000221). 2–4-year-olds differed significantly from 4–6- and and 6–8-year-olds (H _(2)_ = −14.229, p=.009; H _(2)_ = −18.991, p=.00014, Bonferroni corrected, respectively), while object identity accuracy did not show any difference between the three age groups (Figure 3 B).

When looking just at children’s choice of locations and objects, without taking their sequence into consideration, they perform well over chance across all ages (locations: 2–4-year-olds: .69 ± .08; 4–6-year-olds: .90 ± .04; 6–8-year-olds: .97 ± .02 paired two-tailed t-test p= .0000365, p< .00001, p< .000001, respectively; objects: 2–4-year-olds: .67± .06; 4–6-year-olds: .70± .05; 6–8-year-olds: 1± .00 paired two-tailed t-test p= .000065, p= .00005, p= .000000, respectively) indicating that young children do not have difficulty remember the objects and places themselves (Figure 3 B, gray lines). Nevertheless, there is significant improvement in both spatial location and object identity over development (Kruskal Wallis one-way ANOVA, H _(2)_ =16.835, p=.000221; H _(2)_ =28.696, p=.0000001, respectively), with 2–4 performing worse than both 4–6 and 6–8 year-olds in spatial location (H _(2)_ =-14.229, p=.009, H _(2)_ =-18.991, p=.000012, respectively, Bonferroni-corrected), and 6–8 performing worse than both 2–4 and 4–6 year-olds in object recognition (H _(2)_ =-24.750, p=.0001, H _(2)_ =-21.188, p=.000021, respectively, Bonferroni-corrected). In order to address the possibility that the early development of space-time binding could simply be an artifact of an improvement in children’s spatial memory (and not in its temporal organization), a GZLM was performed using space-time performance as the dependent variable, age-groups as the independent factor and spatial location accuracy as a covariate (Figure *2B left*). Performance in the Space-Time test varied significantly between the three age-groups (F_(2)_ =6.755, p=.002). 2–4-year-olds (.48 ± .07) differed significantly from 4–6- and 6–8-year-olds in space-time binding adjusted for spatial location accuracy (.74 ± .05, p=.014; .82 ± .05, p=.002, respectively, Bonferroni-corrected) suggesting that the variance in the model is not explained simply by the change in spatial memory, but by the developmental changes in the binding of space and time. Similar analyses were performend on the Object-Time Test (Figure 3 B, right), using object-time binding performance as the dependent variable, age-groups as the factor and object identity accuracy as a covariate. The overall statistically significant difference in the Object-Time Test between the age groups was no longer significant once object-time binding performance was adjusted for object identity accuracy (F_(2)_ =.871, p=.423), suggesting that the age effect in object-time binding is mostly driven by object recognition capacities. All together, these results suggest that the development of temporally continuous representation of locations does not simply rely on spatial memory capacities but on the improvement in space-time binding processes specifically, and that this pattern in development is not true of remembering a sequence of objects.

### Predictive Role of space-time binding for EM representation

In order to investigate whether space-time binding processes may predict the EM development, we performed an ordinal regression analysis using performance in Space-Time and Object-Time Tests as predictors, and performance in the EM Test as dependent variable, with age as covariate (see Figure 4). All the variables, with the exception of the covariate, have an ordinal distribution representing four scores in a range between .00 (“Not Accurate”) and 1.00 (“Strongly Accurate”). The model was statistically significant (χ^2^(7) = 24.417, *p* = .001), showing a significant relationship between the EM (scored 1.00, as referent value), and the total failure in Space-Time Test (Wald χ^2^ (1) = 4.067, p <.05; negative logit regression: Exp (B)= .137) (the Wald score describes the difference between a maximum likelihood point estimate of a single parameter and a hypothesised value to its standard error). This result means that the failure in Space-Time Test reduces the probability of being strongly accurate in EM Test, and that only space-time, but not object-time binding, is a predictor of the EM accuracy. These results are in line with the literature underlying a scaffolding role of space-time binding for episodic memories, highlighting the ontogenetic emergence of this cognitive process.

**Figure 4.**
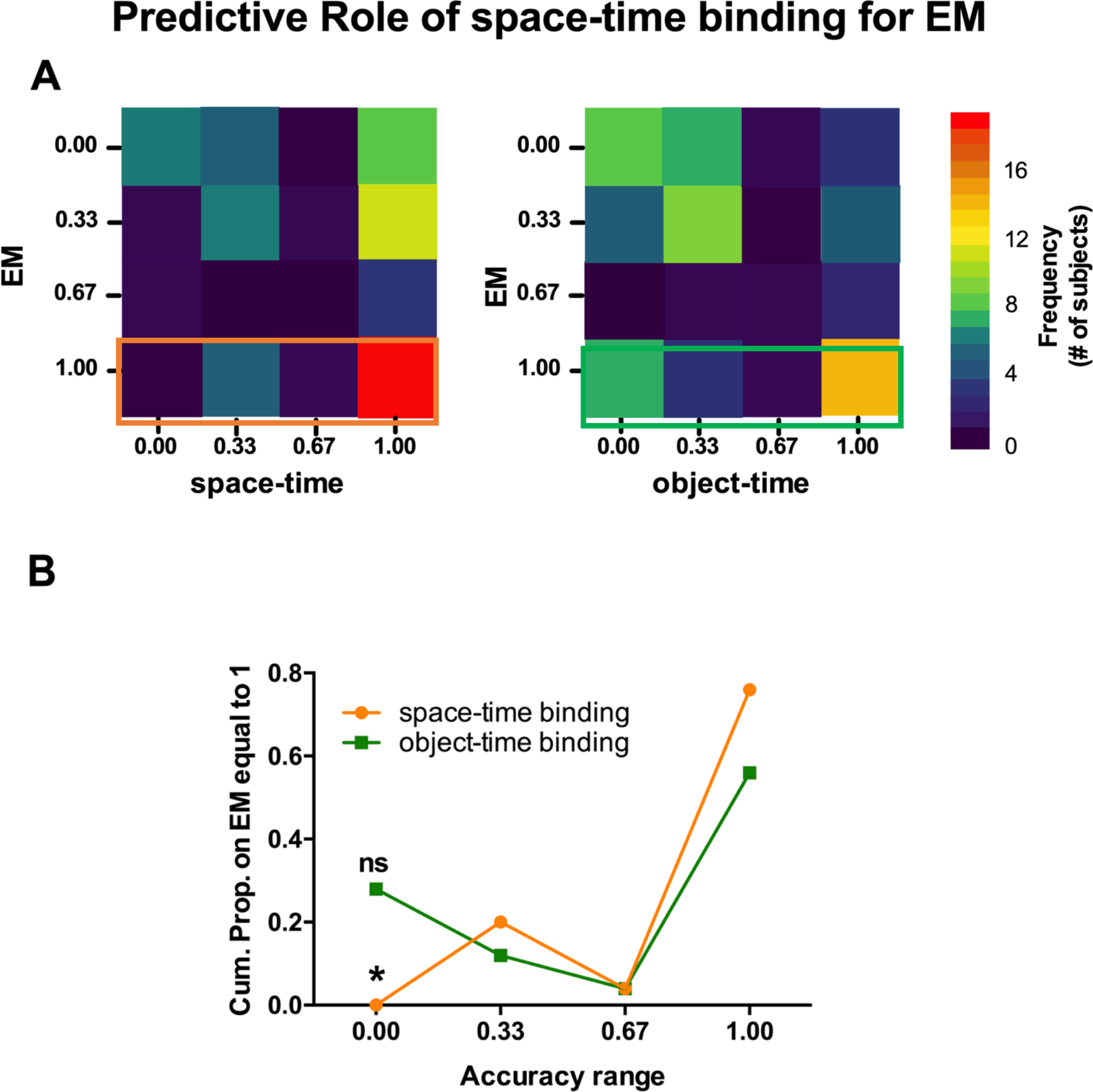
Predictive Role of space-time binding for EM representation **A.** shows the frequencies of the co-occurrences between each EM Test score and Space-Time (right), and Object-Time (left) Tests. **B.** presents the respective cumulative proportions of Space-Time and Object-Time Tests on accurate scores (equal to 1) on the EM Test. On the basis of the Ordinal Regression Model results, when the performance in Space-Time Test is equal to 0.00 (failure in binding space and time), the probability of getting the EM test completely accurate is significantly lower. ns; p < 0.05 *

### Detailed analysis on the EM Test components: enhancement of space-time binding

We performed a detailed analysis on the EM Test components (i.e. spatial location, object identity, space-time, object-time, object-space binding processes) in order to investigate whether space-time binding processes may be a scaffold for the EM emergence. Figure 5 A shows the distribution of each EM Test binding component in the TD sample divided by age groups. Performance on space-time and object-space binding varied significantly by age-group, while object-time did not show any significant effect (Kruskal Wallis one-way ANOVA, *object-time:* H _(2)_ =5.437, p=.066*; space-time*: H _(2)_ =8.678, p=.013; *object-space*: H _(2)_ =12.029, p=.002). 2–4year-olds and 6–8-year-olds differed significantly in space-time and object-space (H _(2)_ = −13.133, p=.040; H _(2)_ = −19.075, p=.003, respectively), while 4–6-year-olds and 6–8-year-olds differed significantly only in space-time (H _(2)_ = −11.769, p=.038). Interestingly, 2–4-year-olds perform as well as 4–6-year-olds children in their ability to bind space and time components in the EM Test. In fact, if we compare these results with those obtained in the Space-Time Test (Figure 5B), we observe that 2–4-year-olds do better in the space-time component of the EM Test than the Space-Time Test (Z=−2.137, p=.033, Wilcoxon signed-ranks test). In contrast, there is no significant difference between the object-time component of the EM Test in comparison with the Object-Time Test (Z=−1.742, p=.082, Wilcoxon signed-ranks test).

**Figure 5.**
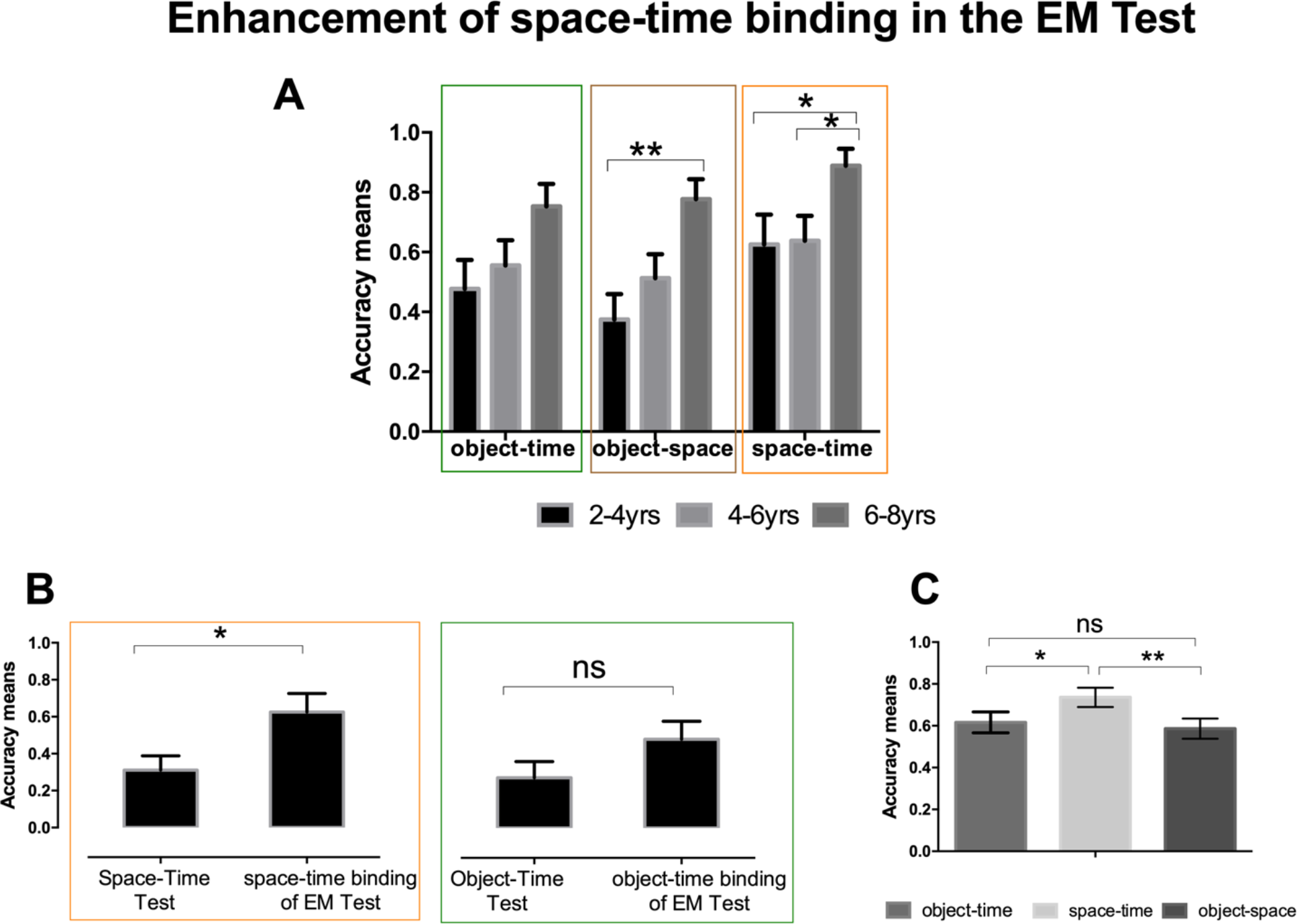
Enhancement of space-time binding in the EM Test *A.* The graphs present the accuracy means for Object-Time, Object-Space, and Space-Time binding components of the EM Test in the TD sample divided by age-groups (2–4; 4–6; 6–8 yrs). *B.* presents the accuracy means of Space-Time Test compared to space-time binding scores of the EM Test, and Object-Time Test compared to object-time binding of the EM Test for 2–4-years old children. *C.* presents the distribution of each EM binding components across all TD subjects. ns; p > 0.05 *; p ≤ 0.05 **

Why are even the youngest children able to bind space and time in the EM Test when the task requires a higher mnemonic load, compared to the Space-Time Test? One possibility may be that because the EM Test was always conducted last, there is a practice effect in terms of understanding the task. However, it is noted that such a practice effect is not found for object-time component. Another possible explanation is that the presence of objects makes the spatiotemporal memory stronger through the association of object content to spatial location. In fact, that may be the very mechanism by which objects get bound onto the spatiotemporal continuum of episodic memory. In Figure 5A, it is possible to see that object-space binding follows a developmental pattern that highly resembles that of object-time. Consistent with this, space-time accuracy is significantly higher than both object-time and object-space accuracy, while object-time and object-space are not different (Z= 2.058, p=.040; Z=2.795, p=.005; Z=-.715, p=.475, respectively, Wilcoxon signed-ranks test). Finally, when we performed a Sperman’s correlations between EM Test and each binding component, we saw that all the variables are strongly correlated to each others (Table 1A). However, taking into consideration the fact that these are related variables, we looked at their partial correlations (Table 1B) and found that if we control for object-space binding, then object-time and space-time binding are no longer significantly correlated (r=.093, p=.456) which suggests that the successful binding of object-time to space-time is mediated by forming associations between object and space. In contrast, if we perform partial correlations controlling separately for object-time and space-time binding (Table 1C), results still remain significant (r=.330, p=.007; r=.566, p=.000001, respectively). In other words, if full EM requires the co-occurrence of remembering where-when with what-when, it seems that the memory of what and where (object-space) plays a contributing role in that process.

**Table 1.**
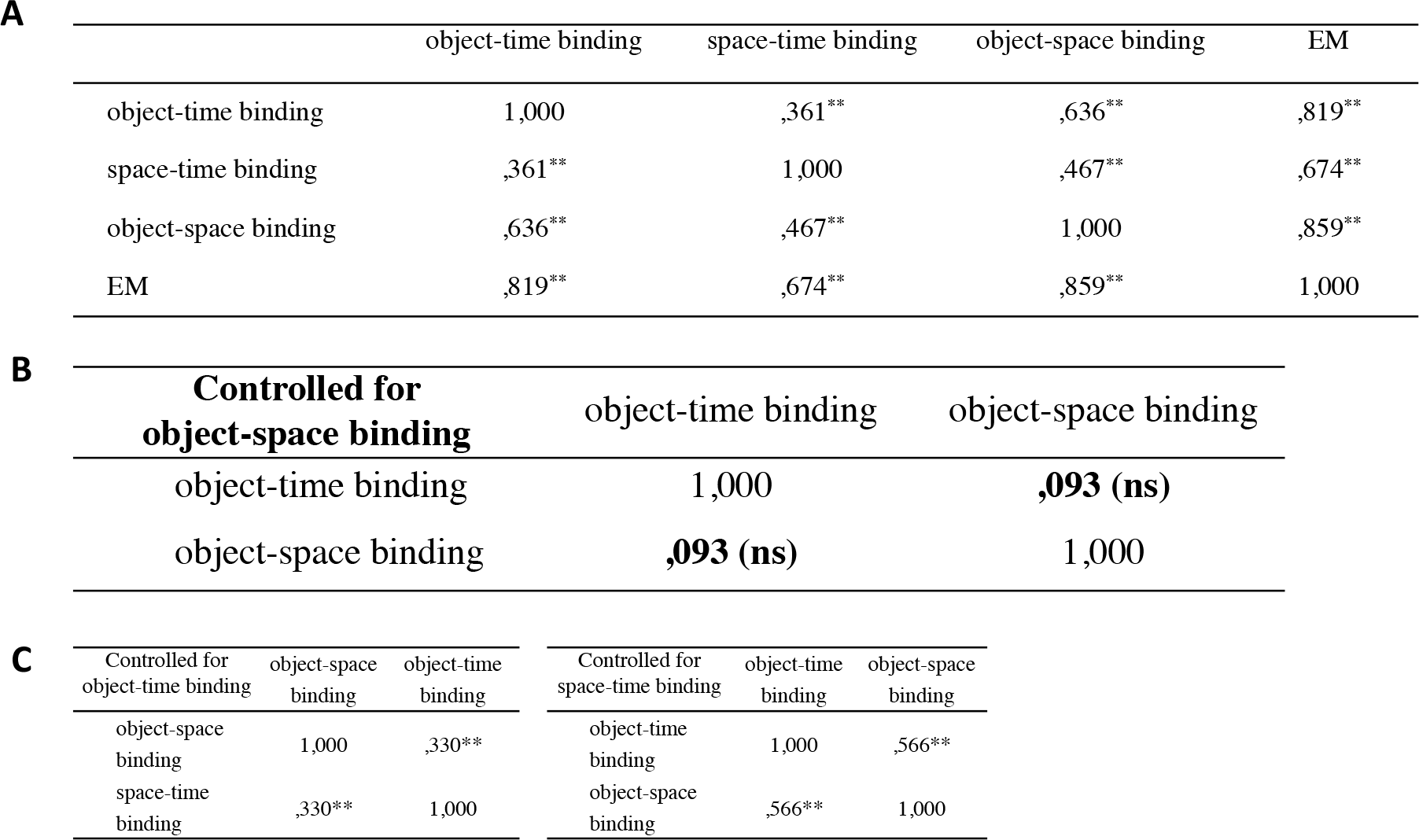
Correlations between EM Test components *A.* Pearson’s correlation for object-time, object-space, and space-time binding components and the total score of the EM Test in the TD sample. *B.* Partial two-tails correlations between object-time and object-space binding components controlled for the effect of object-space binding. *C.* Partial two-tails correlations between object-space and object-time binding components controlled for the effect of object-time binding (*left*), and the correlation between object-time and object-space binding components controlled for the effect of space-time binding (*right*).

|  |  |  |  |  |  |
| --- | --- | --- | --- | --- | --- |
| **A** |  | object-time binding | space-time binding | object-space binding | EM |
|  | object-time binding | 1,000 | ,361** | ,636** | ,819** |
|  | space-time binding | ,361** | 1,000 | ,467** | ,674** |
|  | object-space binding | ,636** | ,467** | 1,000 | ,859** |
|  | EM | ,819** | ,674** | ,859** | 1,000 |

|  |  |  |  |
| --- | --- | --- | --- |
| **B** | **Controlled for**<br>**object-space binding** | object-time binding | object-space binding |
|  | object-time binding | 1,000 | **,093 (ns)** |
|  | object-space binding | **,093 (ns)** | 1,000 |

**Table 1.** Correlations between EM Test components **A.**
|  |  |  |  |  |  |  |
| --- | --- | --- | --- | --- | --- | --- |
| **C** | Controlled for<br>object-time binding | object-space<br>binding | object-time<br>binding | Controlled for<br>space-time binding | object-time<br>binding | object-space<br>binding |
|  | object-space<br>binding | 1,000 | ,330** | object-time<br>binding | 1,000 | ,566** |
|  | space-time<br>binding | ,330** | 1,000 | object-space<br>binding | ,566** | 1,000 |

### Study 2: Atypical development of hippocampal binding components in Williams Syndrome

Williams Syndrome is a rare genetic disorder caused by a microdeletion of about 25 genes in region q11.23 of chromosome 7. WS is a particularly interesting case of inquiry the EM binding processes because of its abnormal hippocampal development. Structurally, the anterior hippocampus is significantly larger than typical controls while the posterior is relatively smaller. However, despite the larger size, the anterior hippocampal include depressed synaptic activity and a bilateral reduction in cerebral blood flow in the hippocampus and entorhinal cortex (Meyer-Lindenberg et al., 2005). WS patients present a unique cognitive profile that includes relative strengths in domains such as language production and face recognition together with severe deficits in a range of spatial functions (Landau & Ferrara, 2013). Previous studies conducted with WS patients have reported impairments in both visuospatial processing (Mervis et al., 2000; Vicari et al., 2005; Martens et al., 2008) and spatial navigation (Lakusta, Dessalegn, & Landau, 2010; Farran et al., 2010; Foti et al., 2011; Mandolesi et al., 2009; Smith et al., 2009; Nardini et al., 2008; Meyer-Lindenberg, Kohn, Mervis, Kippenhan & Olsen, 2004).

Subjects with WS have been reported to have impaired memory for associated pairs of items (Costanzo et al., 2013) and word-location pairs (Edgin, 2010) compared to controls, indicating a memory binding deficit in this population. While there is a large body of literature on spatial memory in WS, no study has specifically investigated the ability to represent temporally continuous memories in this syndrome. Therefore, Study 2 tested WS subjects (compared to TD controls) in the same task described above in Study 1, to selectively assess single EM components and their binding with time, allowing us to speculate on the role of the hippocampus for episodic memory and even its potential genetic basis.

## Results

### Impairment in space-time binding and its impact on EM capacities in WS

We investigated memory binding processes in WS subjects, comparing their performance with those of mental (MA) and chronological aged (CA) controls (Figure 6A). Performance on the three tests significantly differed across groups (Independent-sample Kruskal Wallis 1-way ANOVA: Object-Time Test: H (2) =6.823, p=.033; Space-Time Test: H (2) =14.658, p=.001; EM Test: H (2) =23.004, p=.000010). WS participants performed significantly worse compared to MA controls in space-time binding and in EM (H (2) =−10.409, p=.028; H (2) =−13.568, p=.022, respectively, Bonferroni-corrected), while their memory for ordered recall of items was only delayed compared to CA (H (2) =−11.591, p=.040; Figure 5A). Interstingly, spatial location and object identity accuracy were both highly accurate and did not show any significant difference across the three groups (Figure 6B).

**Figure 6.**
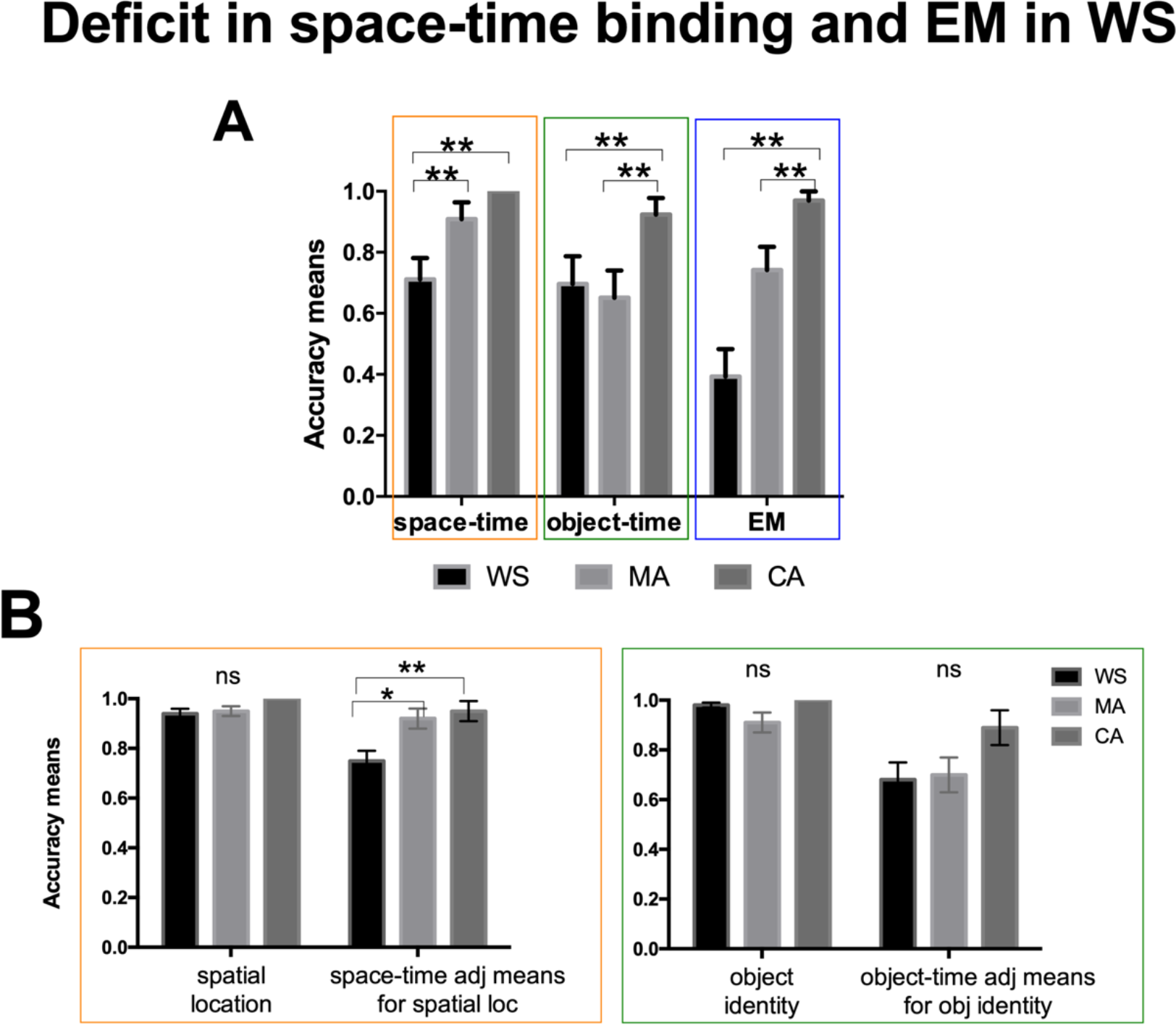
Development of binding components in WS patients *A.* The graphs present the accuracy means for Space-Time, Object-Time and EM Tests in WS patients, compared to MA and CA controls. *B.* The graphs present the means for spatial location accuracy, and space-time binding adjusted for spatial location accuracy (left), and for object identity accuracy, and object-time binding adjusted for object identity accuracy (right). Adjusted means were calculated using GZLM. ns; p > 0.05 *; p ≤ 0.05 **

In order to address the possibility that the failure in space-time binding could be driven simply by a deficit in WS subjects’ spatial memory accuracy (and not its relation to temporal continuity), we performed a GZLM using performance in Space-Time Test as dependent variable, groups (WS, MA, CA) as factors and spatial location accuracy as a covariate (Figure 6 B *left).* Performance in Space-Time Test differend significantly between the three groups having adjusted for spatial accuracy (F_(2)_ =6.927, p=.002), with WS subjects (.75 ± .04) performing significantly worse than MA and CA controls (.92 ± .04, p=.010; .95 ± .04, p=.004, respectively, Bonferroni-corrected). Perhaps, the variability in the model is not driven by the variability in spatial accuracy, but by those in space-time binding. A similar analysis was conducted using performance on Object-Time Test as dependent variable, groups as factors and object identity accuracy as covariate (Figure 6 B *right)* showing that the overall statistically significant difference in object-time binding between different groups disappears once its score has been adjusted for the covariate (F _(2)_ =2.299, p=.109). Perhaps, the group effect in Object-Time Test is mostly driven by object identity recognition capacities. All together, these results suggest that the deficit in space-time binding in WS subjects does not simply rely on spatial memory difficulties but on the binding processes deficits. As was the case for typically developing children in Study 1 (see Figure 3 B), temporal information was particularly relevant when the task requires to recall a sequence of locations rather than objects.

In order to investigate whether the WS performance may be predicted by the general cognitive abilities of participants, we performed Spearman’s Rank Order correlations between WS subjects’ mental age, performance in Space-Time, Object-Time Tests, spatial location and object identity accuracy (Figure 7). A significant correlation was found only between mental age and spatial location accuracy (r=.54, p=.003), but not between mental age and space-time binding (r=.24, p=.355). This means that as mental age (and general intelligence) increases in WS, the capacity to remember location improved; however, space-time binding deficit is specific and is not related to the general cognitive abilities of WS patients.

**Figure 7.**
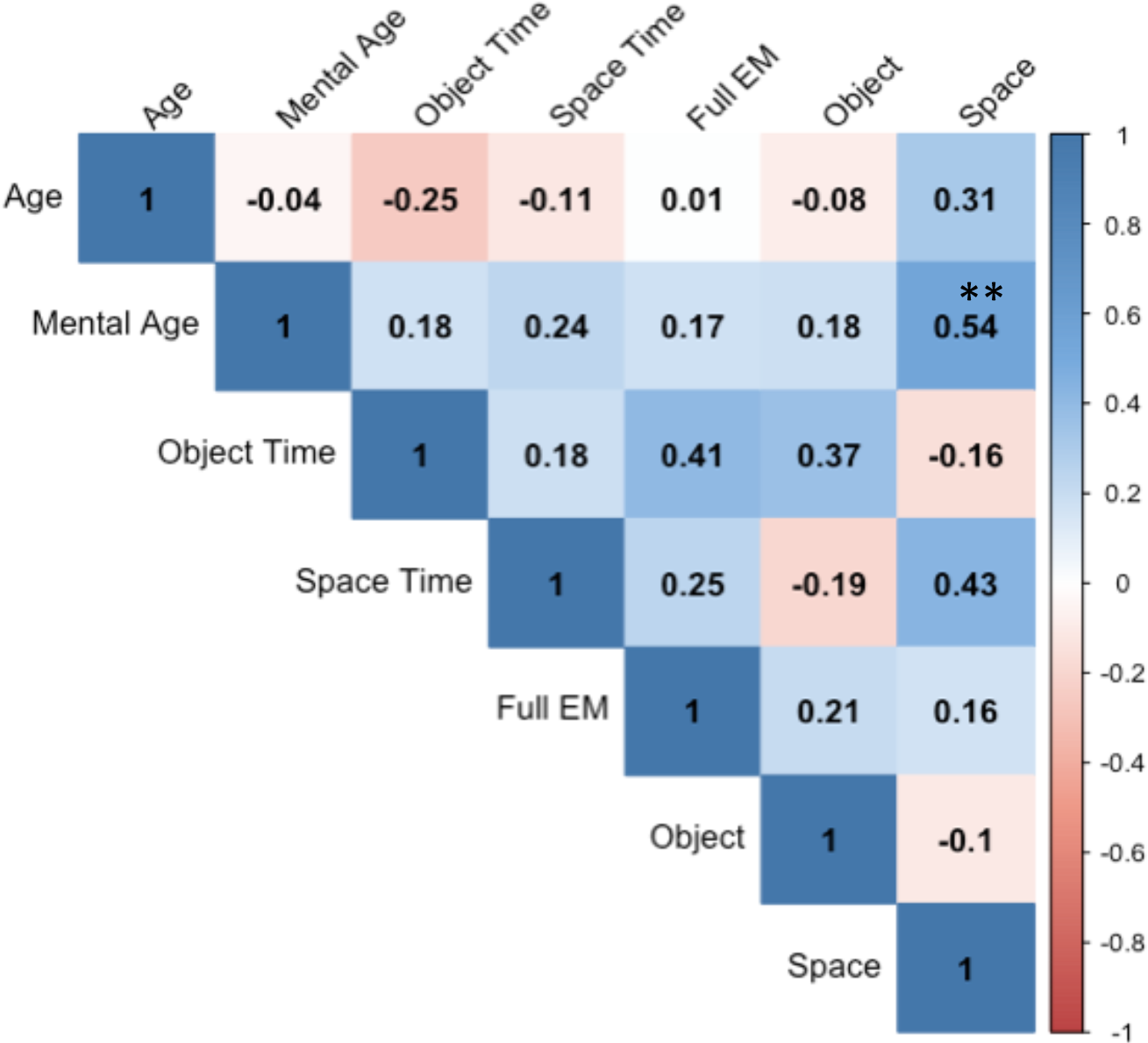
The graph presents Spearman’s Rank Order correlations performed between WS mental age, and the different single and binding components. A significant correlation emerges between mental age and spatial location accuracy (p<<.005) but not between mental age and the other components. ns p > 0.05; *p ≤ 0.05; **p ≤ 0.01; ***p ≤ 0.001.

## Discussion

### Typical Development of EM binding components

In Study 1, we set out to investigate whether space-time binding is a scaffold for the EM representations to be built upon. We found that the ability to fully bind the what, when, and where components of an event develops around 6 years of age, consistent with other studies of episodic memory in children (Sluzenski et al. 2006; Lloyd et al., 2009). Additionally, however, we showed that the emergence of episodic recall is preceded by the development of a competence in coherently binding together spatial and temporal information from 2 to 4 years of age and that this particular memory-binding component predicts EM ability across individuals. Moreover, children’s ability to embed objects with time is significantly correlated with their ability to bind those objects to space, which may in effect provide a spatiotemporal framework (i.e., we are able to remember the sequence of object because we are able to remember the sequence of spaces) necessary for successful EM.

The interpretation that the association of object content to spatial context is what allows objects to be embedded in episodic memory is supported by direct recordings of neuronal activity in the human hippocampus from subjects performing computer-based navigation tasks (Miller et al., 2013) and is consistent with theories that posit that the spatiotemporal context functions as a primary scaffold for memories of events (Ekstrom & Ranganath, 2017). However, our results add to that view the ontogenetic origins of the basis for such EM representations in early human development. Our findings are also consistent with previous studies in cognitive development indicating a significant improvement in the use of environmental cues in spatial navigation around 5 or 6 years of age (Gianni, de Zorzi, & Lee, 2018; Hermer-Vazquez et al.1999, Lee & Spelke, 2010, Vasilyeva & Lourenco 2012). Improvement in spatial navigation could be related to the emergence of a more mature episodic memory around the same age. Lambert et al. (2015) suggest that the development of two shared mechanisms underlie both episodic memory and allocentric navigation: pattern separation, necessary to distinguish different memories and supported by the dentate gyrus (Ribordy et al., 2013), and pattern completion, needed to integrate the elements of a memory and supported by CA3 and CA1 (Hunsaker & Kesner, 2013; Hindy, Ng, Turk-Browne, 2016). Protracted maturation of the dentate gyrus (Jabès et al., 2010) as well as its connections with CA3 (Clelland et al., 2009) might underlie the creations of allocentric representations of space and the emergence of episodic recall.

### Atypical development of episodic memory: Williams Syndrome

In Study 2, we investigated how the emergence of memory-binding abilities is affected by hippocampal abnormalities in WS. We showed that WS participants, unlike 4-year-olds TD children, struggle to recall space-time binding information and perform even worse when attempting to bind object contents into their spatiotemporal context in EM.

Accurate memory for the spatial context of objects has been linked to activity in the anterior hippocampus (DeMaster & Ghetti, 2012; DeMaster et al., 2013), while the posterior hippocampus correlates with navigation expertise (Maguire et al., 2000). Given that the hippocampus of WS individuals exhibits structural and functional abnormalities along the anterior-posterior axis, we might hypothesize that their impairment in spatiotemporal binding may be related to their smaller posterior hippocampus and that their difficulty in further binding elements into the episodic may be related to the decreased activity in the anterior hippocampus and entorhinal cortex.

Interestingly, despite a clear and consistent deficit in spatio-temporal and episodic recall, the spared memory for sequences of objects in WS suggests an alternative strategy of object-related processing that might rely in part on connections with the prefrontal cortex (Naya et al., 2017). Additionally, this object-related success is consistent with spatial navigation studies that show better (Lakusta et al., 2010), qualitatively different (Ferrara & Landau, 2015) featural landmark use in WS compared with typically-developing children.

Finally, it is important to note that memory deficits presented by WS participants as well as by younger TD children, are about binding components of memory, rather than recognizing them in general. In fact, children and WS subjects were highly accurately in picking out the correct places or objects; the just could not place them in a continuous sequence. This finding is consistent with past work that shows a dissociation between recognition processes and memory-binding in TD children (Picard et al., 2012, Sluzenski et al., 2006, Lloyd et al., 2009, Hayne & Imuta, 2011; Lorsbach & Reimer, 2005) and a preserved object recognition in WS (Costanzo et al., 2013; Vicari et al., 2005).

## Conclusions

Time seems to have a special relation to space in that the first component of episode-like memory to emerge in development and be impaired in WS is the combination of spatial and temporal information. Not only that, time seems to depend on space in order to bind together conceptual content such as object identity into its folds. These findings allow us to speculate on a potential layered organization of the episodic system in which the hippocampus acts as a hub of information (Battaglia et al., 2011; Diana et al., 2007; Gotgay et al., 2006) but that which prioritizes spatiotemporal binding that makes mental time travel possible.

Further progress in our understanding of how episodic memory develops in children and how they become impaired with hippocampal abnormality is crucial for several reasons. First, the neural mechanisms underlying EM has been implicated not only in memory but in imagination or future mental time-travel (Maguire and Mullally, 2013; Schurr et al., 2018). Understanding how EM performance can change will indirectly shed light on the development (and impairment) of high-level cognitive abilities such as creativity and imagination. Second, it has been proposed that the hippocampus adaptively organizes initially-encoded episodic memories into conceptual knowledge that drives novel behavior (Constantinescu, O’Reilly, Behrens, 2016; Mack, Love, Preston, 2018). Insight into the origins of hippocampus-dependent memory binding mechanisms may help characterize how more abstract hippocampal maps beyond just spatiotemporal ones, develop and form in human cognition. Finally, we believe that there is an urgent need for a better characterization of the nature of episodic memory processes in order to be equipped with the scientific knowledge to support a society in which we can enjoy long and healthy lives, not just physically but also mentally.

## Methods

### Participants

*Study 1* involved 67 children (34 females) between the ages of 2.52 and 8.74 (M=5.53; SD=1.63). All children were tested at the Cognitive Neuroscience Developmental Lab of CIMeC, University of Trento. No child had any significant medical or psychological deficit. 11 additional children were excluded from the study due to medical problems, prematurity, important language development delays and refusal to follow instructions or carry out the task.

*Study 2* involved 22 Williams Syndrome subjects (11 females). WS participants belonged to two different Italian Williams Syndrome Associations and were tested during two different yearly WS retreats and by appointment at the Clinical Genetic Unit, Fondazione IRCCS Ca Granda Ospedale Maggiore Policlinico in Milan. Importantly, in each location we recreated the same experimental environment in terms of room size and landmark positions. Each WS participant was individually matched for mental and chronological age, gender and trial order with typically developing controls. WS subjects ranged in chronological age from 12.86 to 43.78 (M = 24.84; SD = 8.11) and mental age range: 5.11–8.96 (M = 6.44; SD =0.98). For a measure of non-verbal mental age, WS participants were administered the four subscales of the Brief IQ Visualization and Reasoning battery of the Leiter International Performance Scale Revised (Leiter-R, Roid & Miller, 1997).

### Procedure

Participants were tested on a non-verbal object-placement task developed to allow isolated testing of the single components that make up an episodic memory, namely object, location and their temporal organization. Each experimental session was composed of three tests (Figure 1): The Space-Time Test was designed to measure the ability to remember spatial locations and organize them in time but did not require object recognition. The Object-Time Test measured the ability to remember different objects in a temporal order, while keeping spatial location constant. Finally, The EM Test assessed the ability to combine object identity, spatial locations, and their temporal order together into one representation. All subjects completed all three tests. The Space-Time Test and Object-Time Test were counterbalanced in their order, and then the EM Test was always presented last. The selected objects, locations, and temporal sequences were always randomized across subjects and within trials. The task was straightforward: the experimenter sequentially placed three objects in boxes laid out in the experimental room while the participant was watching (demonstration phase), and then the participant was asked to do the same thing as the experimenter by placing the objects themselves in order to provide the participant of a controlled episode involving himself/herself (encoding phase). Finally, after a delay (verbal interference task), the participant was asked to re-enact his own previous actions (retrieval phase).

Before beginning the experiment, two training tasks were administered in order to elicit imitation behavior in children. In the first one, the experimenter hid one of three colored crayons in one of three grey plastic cups, while in the second one he produced some motor sequences while standing up and facing the participant. The participant was then asked to repeat the experimenter’s behavior. If the participant struggled to understand or failed to reproduce the correct behavior, the experimenter repeated the sequence while emphasizing the relevant information. The training ended after the child had successfully copied the experimenter twice.

To start the experiment, the experimenter brought the participant to the starting position located at one end of a rectangular room, in front of a large door. Once the participant was standing in front of the door facing the room, the experimenter proceeded to give the instructions by showing an animal finger puppet (e.g., a lion) and telling the following short story: “You know, this little lion loves cookies; she always bakes too many and then has to hide them. Now, pay attention carefully to how I hide them, because afterwards you will have to do it just like me. Otherwise she’s not going to find them later!”. The participant was then shown the plastic container with the specific “cookies” (objects) that the animal had “baked” and had five seconds to observe them or manipulate them. The container was then placed on a close table, out of view from the child. Then the participant watched the experimenter hide three objects into a subset of the five boxes that were distributed across the room. For the Space-Time Test, children were provided with three identical objects that they had to place in three of the five boxes in the room. For the Object-Time Test, children had to choose three of five different objects to place into a single box nearby. For the EM Test, children had to choose three of five different objects, and place them into three of five different boxes. The objects were always hidden one at the time: the experimenter would pick up one, show it to the child for a couple seconds, then proceed to hide it, and then return to the starting point. After each of the three objects were placed in a box, the child and the experimenter led the child to turn around to face away from the boxes under the pretense of hiding from strong winds, while another experimenter collected the objects.

For the participant to encode his/her own episodic experience, the encoding phase involved the participant being presented with the same objects and instructed to “hide the cookies” just like the experimenter did. If the participant attempted to pick up more than one object at once, the experimenter would remind him/her to hide them one at the time. Once each of the three objects had been hidden, child and experimenter would again face away while they were picked up by a second experimenter. After the encoding episode, a 3-minute interference task was administered. The participant and the experimenter sat down together at a table present in the room under the pretense of teaching the animal puppet a few words. A verbal task was then administered for a total of 3 minutes. Finally, during the retrieval phase, participant and experimenter returned to their starting position by the door and the participant was presented with the objects and asked to “hide the cookies” as s/he had previously done. After each retrieval phase, following unseen retrieval of the objects, the participant was rewarded with a sticker (regardless of his/her performance).

In order to investigate the potential effect of learning a particular trajectory or movement, few subjects were tested in a “gated condition” during which a plastic gate was introduced right in front of the child, between the encoding and retrieval phases. In order to perform the task, the child was required to walk around the gate, making a different route than the one previously done during the encoding phase. There were no differences between children in this condition, compared to the rest (Space-Time Test *U*= 9.500 p=.548; Object-Time Test *U*= 5.500 p=.151; EM *U*= 8.000 p=.421, Mann-Whitney U Test).

### Scoring

By comparing the subject’s accuracy between the encoding and retrieval phases, we calculated the accuracy in the participant’s memory for the single elements (object identity or spatial location) and the extent to which they were able to bind them in a temporal sequence (object-time binding; space-time binding; EM). For the individual element analysis, the space location and object identity accuracy were calculated giving 1 point for each correct remembered location or object, respectively. For temporal binding, three measures were calculated: space-time binding, giving 1 point for each correctly-remembered location in the correct place in the sequence; object-time binding, assigning 1 point for each correctly-remembered object in its correct temporal sequence; EM, giving 1 point for each different object that is correctly remembered in its correct spatial *and* temporal position (general EM score). The general EM score was decomposed into three further component scores representing space-time, object-time, and object-space binding. For simplicity, each index was calculated on the basis of a proportion of correct scores (maximum raw score being 3).

## Data availability statement

The datasets generated during and/or analysed during the current study are available from the corresponding author on reasonable request.

## Bioethics policy statement

This study was conducted in accordance with the guidelines of the Human Research Ethics Committee of the University of Trento (Italy). Informed consent was obtained from both the participants and their parents or legal guardians.

## Author contributions

M.M. and S.A.L. conceived and designed the study; M.M., N.B and M.F.B. organized subject recruitment; M.F.B performed clinical duties associated with Williams Syndrome subjects; M.M. and N.B. performed data collection; M.M. and N.B. analyzed the data; M.M. and S.A.L. drafted the manuscript.

